# Trascriptome meta-analysis of microalga *Dunaliella tertiolecta* under stress condition

**DOI:** 10.1101/2022.04.11.487921

**Authors:** Bahman Panahi, Mohammad Farhadian, Seyyed Abolghasem Mohammadi, Mohammad Amin Hejazi

## Abstract

Microalgae are photosynthetic organisms, which are considered as a potential source for sustainable metabolite production. Furthermore, stress conditions can affect metabolite production. In this study, a meta-analysis of RNA-seq experiments was performed to evaluate the response of metabolite biosynthesis pathways in *Dunaliella tertiolecta* to abiotic stress conditions, including high light, nitrogen deficiency, and high salinity. The results indicated down-regulation of light reaction, photorespiration, tetrapyrrole, and lipid-related pathways in salt stress. In comparison to salt stress, nitrogen deficiency mostly induced light reaction and photorespiration metabolisms. The up-regulation of phosphoenolpyruvate carboxylase, phosphoglucose isomerase, bisphosphoglycerate mutase, and glucose-6-phosphate-1-dehydrogenase (involved in central carbon metabolism) was observed under salt, high light, and nitrogen stress conditions. Interestingly, the results indicated that the meta-genes (i.e., modules of genes strongly correlated) tended to be located in a hub of stress-specific PPI (Protein-Protein Interaction) networks. Module enrichment of meta-genes PPI networks highlighted the cross talk between photosynthesis, fatty acids, starch, and sucrose metabolism under multiple stress conditions. Moreover, it was observed that the coordinated expression of the tetrapyrrole intermediated with meta-genes involved in starch biosynthesis. The results of the present study also showed that some pathways such as vitamin B6 metabolism, methane metabolism, ribosome biogenesis, and folate biosynthesis responded to different stress factors specifically. In conclusion, the results of this study revealed the main pathways underlying the abiotic stress responses for optimized metabolite production by the microalga *Dunaliella* in future studies. PRISMA check list was also included in the study.

## Introduction

Photosynthetic microalgae have been considered as a potential source of secondary metabolites and proposed as a promising cell factory since they have efficient photosynthesis apparatus and biomass production (Spolaore et al. 2006, Del Campo et al. 2007). Environmental stress conditions can redirect microalga energy flux towards the production and accumulation of fatty acids, starch, and carotenoids (Dellomonaco et al. 2010). This feature is exploited to produce biofuels, pigments, and starch; however, it can reduce metabolite production as well. The growth rates and productivity of microalgae are dramatically diminished under stress conditions (Adams et al. 2013). Growth media optimization is proposed as an effective method to overcome this problem (Feng et al. 2011, San Pedro et al. 2013); however, previous attempts to gain sufficient levels of productivity have failed because the underlying molecular mechanisms are not well understood (Courchesne et al. 2009, Prochnik et al. 2010).

It has been documented that abiotic stresses increase the transcription and activity of key enzymes involved in various metabolite biosynthesis pathways (Ho et al. 2017). Salt stress has also been reported to enhance the activity of glycerol metabolism enzymes such as glycerol-3-*salina* (Breuer et al. 2013). Recent study have also showed that mRNA level of zeaxanthin epoxidase, an enzyme involved in the accumulation and conversion of zeaxanthin, increases rapidly in response to the increased salinity or light/dark cycle (Kim et al. 2019). Similar results were also obtained for the enzymatic activities of fructose-bisphosphate aldolase (FBPA) involved in starch metabolism (Klok et al. 2013). The high light condition phosphate phosphatase, glycerol 2-dehydrogenase, and dihydroxyacetone kinase in *Dunaliella* leads to energy dissipation and the inhibition of photosynthetic apparatus and generates reactive oxygen species (ROS) (Jin *et al*. 2003). Cells can utilize the higher light energy for growth only after adjusting the photosynthetic apparatus and cellular processes (Nymark et al. 2009). The expression of most photosynthesis genes decreases under high light conditions, indicating excess light capture (Binte Safie et al. 2018). Fang *et al*. (2012) demonstrated that about 15% of energy metabolism and carbon metabolism and 25% of Tricarboxylic acid (TCA) related genes were up-regulated under high light conditions. It has also been confirmed that the overexpression of genes involved in carbon metabolism (e.g., fructose -1, 6-bisphosphatase) improves photosynthetic performance as well as the total organic carbon and glycerol production of *Dunaliella bardawil* (Fang *et al*. 2012). The metabolome reprogramming effects of nitrate deficiency and/or high light in *D. tertiolecta* cells has also been examined in the literature. It is revealed that such stresses increase saturated fatty acids and decrease polyunsatured fatty acids (Lee *et al*. 2014).

Despite the broad scientific interest in identifying the responses of *D. tertiolecta* to environmental stresses, the involved biochemical pathways and the underlying genetic mechanisms have still remained unknown (Binte Safie *et al*. 2018). Moreover, contradictory information on the transcriptional regulation of *Dunaliella* under stress conditions has been reported. For example, some studies reported that Glycerol-3-phosphate dehydrogenase *GPDH-c* transcription levels were increased under salinity stress (Kim *et al*. 2010), whereas another study showed that the *GPDH-c* expression level was decreased under salt stress (Alkayal *et al*. 2010). Additionally, various studies have reported different degrees of FBPA expression under abiotic conditions (Cai *et al*. 2013, Kim *et al*. 2010). On the other hand, not all Dunaliella species respond in the same manner to abiotic stress conditions (Kim et al. 2010, Pereira and Otero. 2019).

Transcriptome meta-analysis is the most promising strategies to overcome the abovementioned challenges (Zhang *et al*. 2020). It is an efficient tool for the identification of fundamental and principal genes/pathways and often forms the basis for novel hypotheses (Zhang et al. 2018). For RNA-seq meta-analysis, the p-value combination technique is proposed based on the inverse normalization and Fisher’s methods (Rau A. et al. 2014a). This approach has been successfully applied to gain insight into crop responses to environmental stresses (Rest et al. 2016). More recently, salt stress transcriptome responses in different rice genotypes were elucidated by using RNA-seq meta-analysis (Kong et al. 2019).

The microalga *Dunaliella* is frequently considered for industrial applications because of its potential to produce high value compounds such as beta-carotene, its high growth rate, tolerance under extreme physiological conditions, lack of cell wall, and low energy requirements in harvesting (Einali et al. 2017, Oren 2010, Rammuni et al. 2019).

In the present study, RNA-seq meta-analysis of *D. tertiolecta* was performed under high light, nitrogen deficiency, and high salinity stress conditions. Then, protein-protein interaction (PPI) network analysis was constructed to extend the possible functions of identified meta-genes in different stress conditions.

## Materials and Methods

### Dataset Collection

RNA-seq raw reads were retrieved from the Gene Expression Omnibus (GEO), Sequence Read Archive (SRA), and European Nucleotide Archive (ENA) databases. Raw reads of samples treated under high salinity/NaCl, high light intensity, and nitrogen deficiency conditions were included in the meta-analysis. The details of included datasets are presented in Table 1.

**Table 1.** Raw data set ID, run accession and read count (beforeand after trinuning).

| Data set ID | stress type | Reference | Data base | Run accession | Library layout | Read Count (before trimming) | Read Count (after trimming) |
| --- | --- | --- | --- | --- | --- | --- | --- |
| SRP052730 | Light and Nitrogen | Shin et al., 2015 | SRA | SRR2120818 | SINGLE | 43,014,956 | 38,345,362 |
|  |  |  |  | SRR2120819 | SINGLE | 43,342,701 | 39,921,921 |
|  |  |  |  | SRR2120820 | SINGLE | 42,219,714 | 40,632,920 |
|  |  |  |  | SRR2120821 | SINGLE | 49,394,621 | 47,982,792 |
| SRP081094 | Nitrogen | Not available | ENA | SRR4011621 | PAIRED | 3,765,251 | 2,946,132 |
|  |  |  |  | SRR4011622 | PAIRED | 3,990,351 | 3,011,629 |
|  |  |  |  | SRR4011623 | PAIRED | 3,501,613 | 2,720,419 |
|  |  |  |  | SRR4011624 | PAIRED | 3,852,588 | 2,940,240 |
|  |  |  |  | SRR4011625 | PAIRED | 3,203,153 | 2,708,372 |
|  |  |  |  | SRR4011626 | PAIRED | 3,303,731 | 2,699,457 |
|  |  |  |  | SRR4011627 | PAIRED | 3,536,396 | 2,947,635 |
|  |  |  |  | SRR4011628 | PAIRED | 4,079,043 | 3,745,201 |
| SRP106620 | Salinity | Safie et al., 2017 | ENA | SRR5520490 | PAIRED | 1,370,286 | 1,056,393 |
|  |  |  |  | SRR5520491 | PAIRED | 1,498,758 | 1,139,345 |
|  |  |  |  | SRR5520492 | PAIRED | 1,395,338 | 1,183,222 |
|  |  |  |  | SRR5520493 | PAIRED | 1,421,445 | 1,103,289 |
|  |  |  |  | SRR5520494 | PAIRED | 944,136 | 826,323 |
|  |  |  |  | SRR5520495 | PAIRED | 2,044,161 | 1,838,644 |
| SRP005987 | Salinity | Yao et al., 2017 | GEO | SRR124253 | SINGLE | 21,798,484 | 19,972,634 |
|  |  |  |  | SRR124254 | SINGLE | 19,855,406 | 18,928,637 |
| SRP003930 | Nitrogen and Salinity | Rismani-Yazdi et al., 2011 | ENA | SRR070439 | SINGLE | 305,673 | 299,374 |
|  |  |  |  | SRR070440 | SINGLE | 333,880 | 300,341 |
|  |  |  |  | SRR070441 | SINGLE | 183,731 | 173,922 |
|  |  |  |  | SRR070442 | SINGLE | 229,538 | 200,231 |
| SRP063050 | Light | Safie et al., 2017 | ENA | SRR6517590 | PAIRED | 1,399,689 | 1,200,384 |
|  |  |  |  | SRR6517591 | PAIRED | 1,522,878 | 1,392,774 |
|  |  |  |  | SRR6517592 | PAIRED | 1,572,853 | 1,234,093 |
|  |  |  |  | SRR6517593 | PAIRED | 1,961,343 | 1,893,674 |
|  |  |  |  | SRR6517594 | PAIRED | 1,839,092 | 1,735,468 |
|  |  |  |  | SRR6517595 | PAIRED | 3,441,148 | 2,992,346 |
|  |  |  |  | SRR5514896 | PAIRED | 3,653,262 | 3,481,926 |
|  |  |  |  | SRR5514897 | PAIRED | 2,410,762 | 2,001,018 |
|  |  |  |  | SRR5514898 | PAIRED | 2,199,299 | 1,972,364 |
|  |  |  |  | SRR5514899 | PAIRED | 2,238,539 | 1,999,233 |
|  |  |  |  | SRR5514900 | PAIRED | 3,017,386 | 2,873,087 |
|  |  |  |  | SRR5514901 | PAIRED | 3,721,648 | 3,200,132 |
|  |  |  |  | SRR5514902 | PAIRED | 3,071,374 | 2,963,692 |
|  |  |  |  | SRR5514903 | PAIRED | 2,768,536 | 2,517,291 |
|  |  |  |  | SRR5514904 | PAIRED | 3,017,778 | 2,904,828 |
|  |  |  |  | SRR5514905 | PAIRED | 2,483,867 | 2,018,310 |
|  |  |  |  | SRR5514906 | PAIRED | 2,184,064 | 1,927,438 |

### Processing RNA-Seq data

Raw FASTQ files of the above-mentioned datasets were quality controlled using FastQC software version 0.11.5. In this step, reads with quality score below 30 were excluded. Subsequently, the high-quality reads were trimmed using Trimmomatic software version 0.32 (Bolger et al. 2014) with the following parameters: LEADING: 30, TRAILING: 3, SLIDINGWINDOW: 4:20, and MINLEN: 45. We first mapped the trimmed reads to the *Dunaliella salina* genome (available at https://www.ncbi.nlm.nih.gov/genome/10713) using TopHat2 software (Trapnell et al. 2009). Since mapping rate was extremely low, de novo assembling with strand-specific mode (—SS_lib_type RF) was performed using Trinity software v2.4.0 (Haas *et al*., 2013). In the next step, the trimmed raw reads were aligned to *de novo* assembled transcripts using Kallisto software v0.44.0 (Bray et al. 2016) with default parameters. The expression of genes was determined as Fragments Per Kilobase Million (FPKM) values per gene, and differential expression of genes between treated and untreated samples was estimated by using the edgeR package (Robinson *et al*. 2010). A significant differential expression was defined as a fold change ≥ |2| and a False Discovery Rate (FDR) corrected p-value ≤ 0.05 (Benjamini and Hochberg 1995). Protein orthology was determined using blastx (cut off E-value of 1.0E-10) of assembled transcript against the *Chlamydomonas reinhardtii* annotation. The top five hits were extracted using in-house python scripts.

### Meta-analysis of differential gene expression

To run the meta-analysis, p-values of differentially expressed genes were combined with two “Fisher’s and inverse” normalization methods complimented in metaRNASeq package (Rau Andrea et al. 2014b). To this end, each stress type dataset was analysed separately using the GLM (Generalized Linear Models). The raw *P*-values for each gene from each of the datasets were combined using Fisher and Stouffer’s methods (inverse normal method). Fisher’s method combines the *P*-values from each experiment with one test statistic defined as:

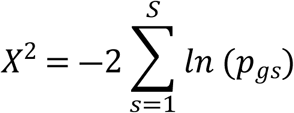

where, *pi* denotes the raw *P-*value obtained from gene *g* in experiment *s. S* is the number of the combined experiments. Under the null hypothesis, the test statistic *χ* follows a χ^2^ distribution with 2 *S* degrees of freedom. This test provides a meta *P*-value, and the classical procedures for multiple testing correction can be applied to obtain the adjusted *P*-values to control the false discovery rate. In the Stouffer’s method, standard normal deviate, the effect size, and its standard are often directly calculated as:

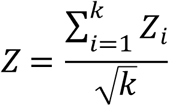

where, *Zi* denotes the one-side *P*-value obtained from gene *i* in the experiment *K* under the null hypothesis indicating no effect in each study. The Benjamini-Hochberg method (Benjamini and Hochberg 1995) was used to correct for multiple testing. Genes with the adjusted meta *P*-value ≤0.05 were considered to be statistically significant.

### Functional analysis of DEGs (Differentially Expressed Genes) and meta-genes

GO enrichment analysis was performed in three categories of biological processes (BP), molecular functions (MF), and cellular components (CC) using the Algal Functional Annotation tool (Lopez *et al*. 2011). *P* ≤ 0.05 was set as the cut-off threshold. To visualize DEGs (Differentially Expressed Genes) and meta-genes within the enriched pathways, MapMan software (Thimm *et al*. 2004) was used. The PPI analysis was also performed to extend the possible functions of meta-genes in the system level. The STRING database (https://string-db.org/) (Szklarczyk *et al*. 2015) was employed for PPI network construction based on the previous knowledge of *Chlamydomonas reinhardtii* at high level of confidence (score ≥ 0.70). Only experimentally validated interactions were included for the network construction. Then the K-means algorithm was used to statistically distinguish functional modules. Finally, the biological functions of each module were enriched using the KEGG database (Kanehisa and Goto 2000). *P*-values were considered to be statistically significant at Benjamini-Hochberg adjusted *P* ≤ 0.05.

## Results and Discussion

### Metabolism overview of differentially expressed genes

The results demonstrated down-regulation of light reactions, photorespiration, tetrapyrrole and lipid-related pathways under salt stress. In comparison to salt stress, nitrogen deficiency mostly induced light reaction and photorespiration metabolisms. Down-regulation of the TCA, starch, and sucrose metabolism-related genes was also revealed under high light stress condition (Fig. 1). It was proposed that the TCA cycle, glycolysis, oxidative pentose phosphate pathway (OPPP), and the carbon concentrating mechanism (CCM) were the main determinants in the carbon flux towards different forms of stored carbon (Tan et al. 2016). Conserved up-regulation of phosphoenolpyruvate carboxylase, phosphoglucose isomerase, bisphosphoglycerate mutase, and glucose-6-phosphate-1-dehydrogenase involved in central carbon metabolism was also observed across high salt, high light, and nitrogen stress conditions (Fig. 1 and Supplementary Table S1). Despite down-regulation of the genes directly involved in the Triacylglycerol (TAG) and Fatty acid (FA) synthesis, up-regulation of glycolytic-related transcripts such as pyruvate kinase and pyruvate dehydrogenase are observed (Fig. 1 and Supplementary table S1).

**Fig. 1:**
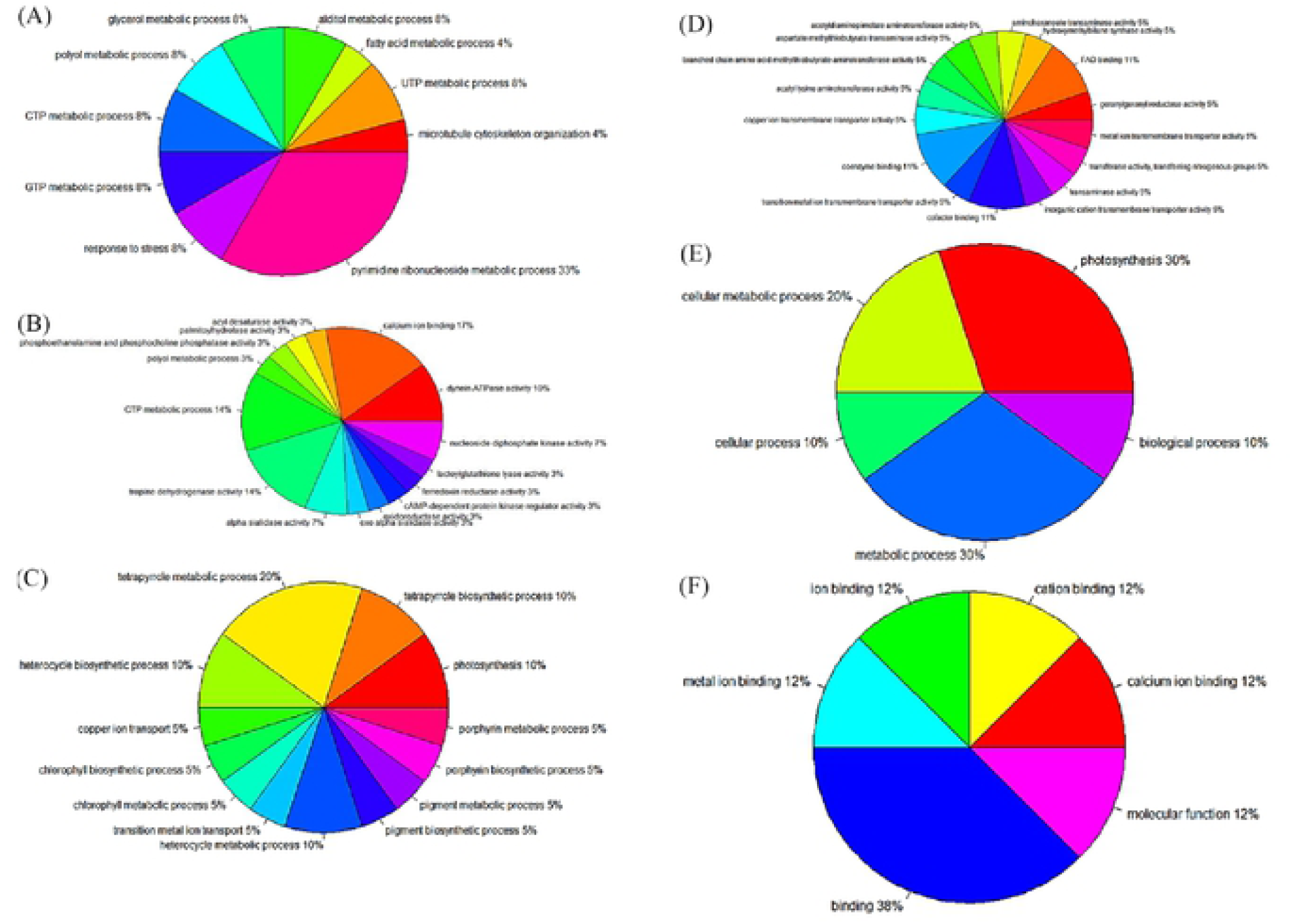
Gen Ontology of DEGs of three Nitrogen stress related experiments. A, C and E, represented the Biological Process, and 8, D and F represented the Molecular functions in the SRP003930, SRP052730 and SRP08!094 experiments, respectively

The findings also demonstrated that nitrogen deficiency down-regulated the ACP, whilst salt stress up-regulated this gene. In response to light stress, the majority of genes involved in lipid metabolism were up-regulated. Ketoacyl ACP synthase and Omega-6 desaturase were two genes with a striking increase in expression (fold change >3) under the same condition. Moreover, in line with the result of this study, it has been reported that genes related to photosynthetic antenna proteins were down-regulated in high light stress condition (Binte Safie et al., 2018). As nitrogen is an important component for the synthesis of chlorophyll and photosystem proteins, the reduction in nitrogen availability could hinder the expression of genes related to photosynthesis (Yao et al., 2017). The results also indicated that nitrogen deficiency up-regulated biotin carboxylase and down-regulated the diacylglycerol kinase. Interestingly, a similar expression pattern of diacylglycerol kinase and biotin carboxylase were observed under both salt and nitrogen stress conditions. Down-regulation of starch degrading enzymes were observed in *D. tertiolecta* cells under high light and nitrogen deficiency conditions (Fig. 1). These results support the idea that *D. tertiolecta* cells tend to accumulate more starch than lipids under high light and nitrogen deficiency conditions (Tan *et al*. 2016).

### Regulation overview of differentially expressed genes

The effects of different stress conditions on regulatory elements are presented in Fig. 2 and Supplementary Figs S4-7. Transcription factors, protein modification, and degradation-related genes, and G-proteins have been frequently identified in the differentially expressed genes. Interestingly, most of the regulatory elements were expressed with stress specific manners. In the category of transcription factors, Cytokinesis related protein (*MOB1*), Histone deacetylase (*HDA*), and C3H zing finger family were down-regulated, whereas Histone acetyltransferase (*HAM*) and Chromatin remodeling factor (*CHR*) were dramatically up-regulated under salt stress condition (Fig. 2). On the other hand, the up-regulation of the *MOB1* and *HDA* was observed under nitrogen deficiency conditions. Moreover, the *CHR* was moderately down-regulated under high light condition. Peptide methionine sulfoxide reductase (*MSRA3*), involved in post-translational modification, was significantly down-regulated under salt stress, whereas it was not involved in the differentially expressed gene list under high light and nitrogen deficiency conditions (Fig. 2). Regarding the protein degradation-related genes, the down-regulation of 26S proteasome regulatory subunit *RPN7*, ubiquitin-activating enzyme E1 (*UBA1*), and active subunits of chloroplast ClpP complex (*CLPP5*) and the up-regulation of ClpD chaperon (*CLPD*) were noticed under the salt stress condition. On the contrary, the *CLPD* was down-regulated under light and nitrogen stresses.

**Fig. 2.**
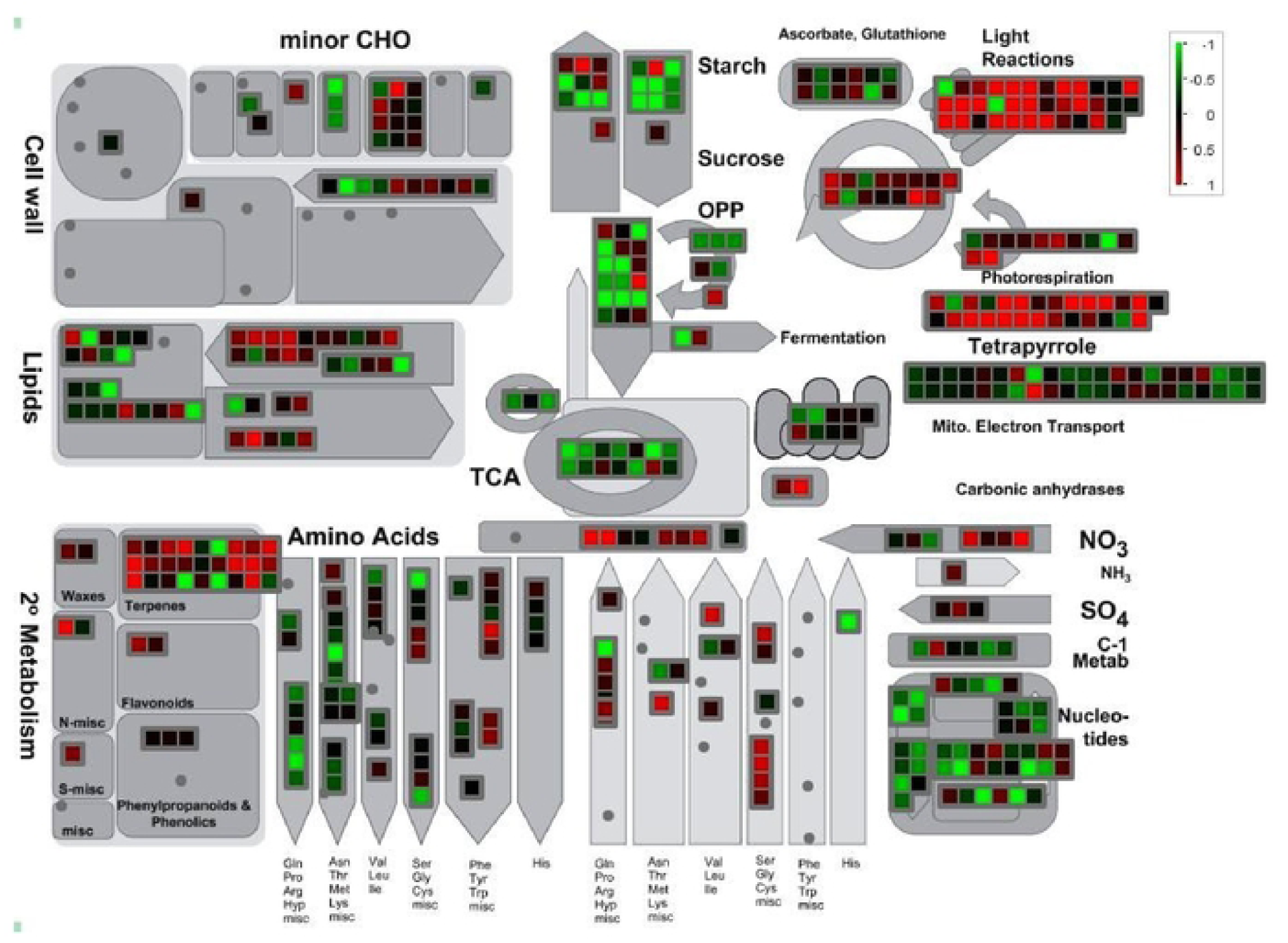
Metabolic overview of differentially expressed genes in Light stress conditions in *D. tertio/ecta*. Green and red box represented the down and up-regulation in corresponding stress, respectively.

*bZIP* (basic leucine zipper) which trigger the cascade of signaling pathways (Zheng et al. 2016), is other key regulatory element of stress responses in *D. tertiolecta*. Results of our study showed that there were similar expression patterns of *bZIP* and *bHLH* (basic helix-loop-helix) (Supplementary Figs. S5-7) with 6-phosphogluconate dehydrogenase (*6PGD*) and glucose-6-phosphate dehydrogenase (*G6PD*, which are involved in oxidative pentose phosphate (OPPP) pathway. The results of our analysis also revealed that *PHD*- and *SET*-transcriptional regulator family is another important regulatory factor in multiple stress responses. As reported, these TFs are regulatory factors in CCM metabolism (in the TCA pathway) and chromatin structure (Bienz 2006). Similar expression patterns of PHD-finger family and some TCA pathway genes (Fig. 2) also highlight this hypothesis.

### Meta-analysis of transcriptome data

Although the transcriptome analysis simplifies some complexities of stress response pathways, it poses some questions in this regard. What are the principal pathways mediating the responses to each type of stress? Are there any putative and unique pathways between different stresses? To answer these questions, we performed the statistical meta-analysis along with the system-level survey of meta-genes. Fig. 3 presents the statistical results of the transcriptome meta-analysis of multiple stresses. Under salt stress condition, 477 (317 down-regulated and 144 up-regulated genes) and 144 (101 down-regulated and 43 up-regulated genes) genes were defined as meta-genes based on the Fisher’s and reverse normalization methods, respectively (Fig. 3A). Under nitrogen deficiency condition, 50 (39 up-regulated and 11 down-regulated genes) and 15 (10 genes up-regulated and 5 genes down-regulated) genes were identified as meta-genes based on the Fisher’s and reverse normalization methods, respectively (Fig. 3B). Moreover, under high light stress condition, 108 (50 up-regulated and 58 down-regulated genes) and 41 (22 up-regulated and 19 down-regulated genes) genes were defined using Fisher’s and reverse normalization methods, respectively (Fig. 3C). Supplementary Table S2 lists the meta-genes for each abiotic stress. Oxygen-evolving enhancer protein (*PSBQ*), superoxide dismutase (*FSD1*) and ascorbate peroxidase (ASP) are examples of the key genes under nitrogen deficiency condition. *FAT1*, encoding the acyl carrier protein thioesterase, and *PPX1*, encoding the protoporphyrinogen oxidase specifically, were determined as the key genes under salinity stress condition (Supplementary Table S2).

**Fig. 3:**
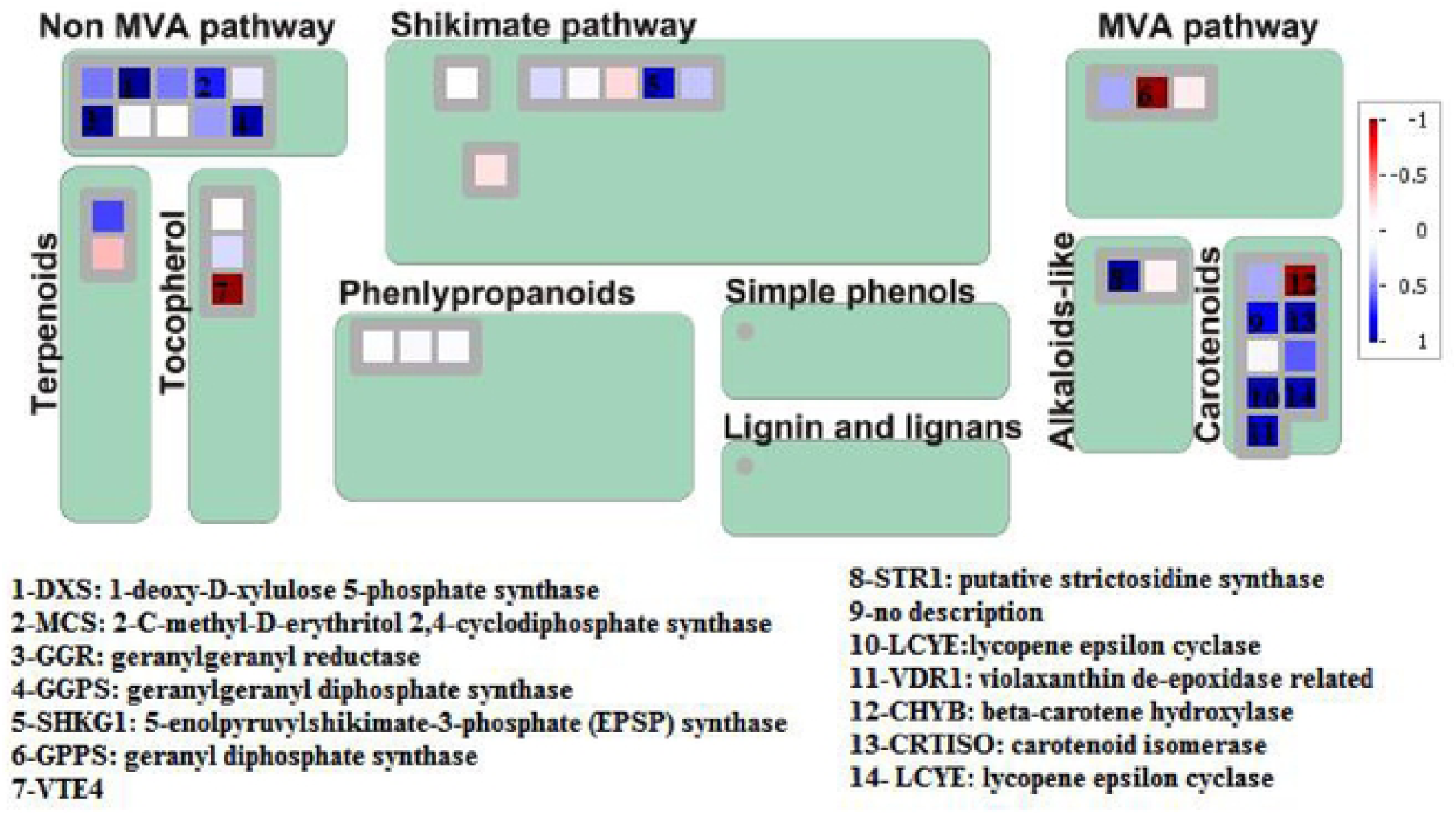
Secondary metabolism genes affected by high light stress. The blue and red boxes represented the up and down regulation severity, respectively.

Interestingly, the results indicate that the key meta-analysis genes tend to be located in a hub situation of the stress-specific PPI networks (indicated by a circle, Fig. 4). Surprisingly, some of these hubs such as serine hydroxyl methyltransferase (*SHMT1*) and *PSBO* are the key regulators of CCM pathways (Fang *et al*. 2017). Not unexpectedly, it the overproduction of storage metabolites seems to be interconnected with defense-related pathways. It is also reported that the functions of *SHMT1* influence resistance to different stress, and the mutation of *SHMT1* leads to increased cell damage due to accumulation of H2O2 (Moreno *et al*. 2005). *STA1*, which is the key enzyme in starch granulation under stress condition, was defined as a core responsive gene (Supplementary Table S2). It has been proposed that the *STA1* mutants significantly reduce granular starch deposition (Wattebled *et al*. 2003).

**Fig. 4:**
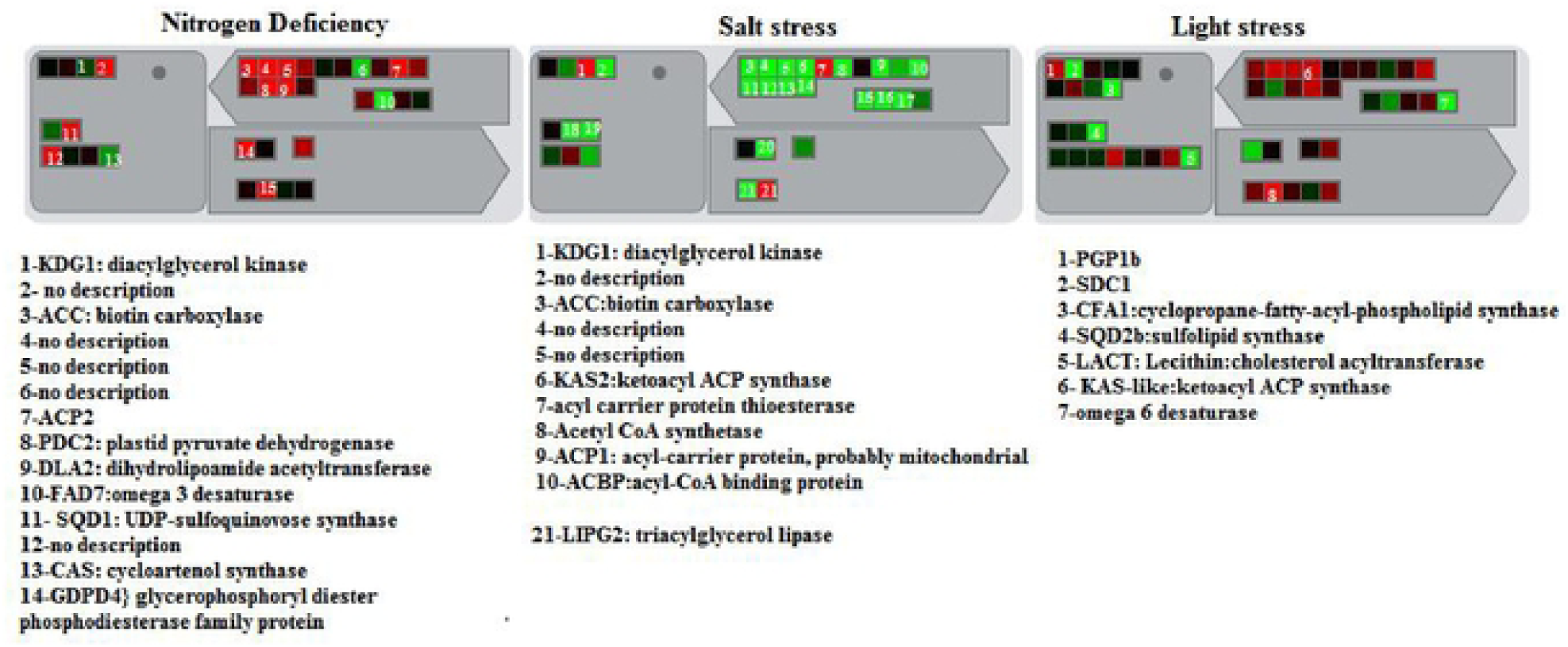
Transcriptional changes of lipid metabolism related genes 1n responses to nitrogen deficiency, salt and high light stress conditions.

**Fig. 5:**
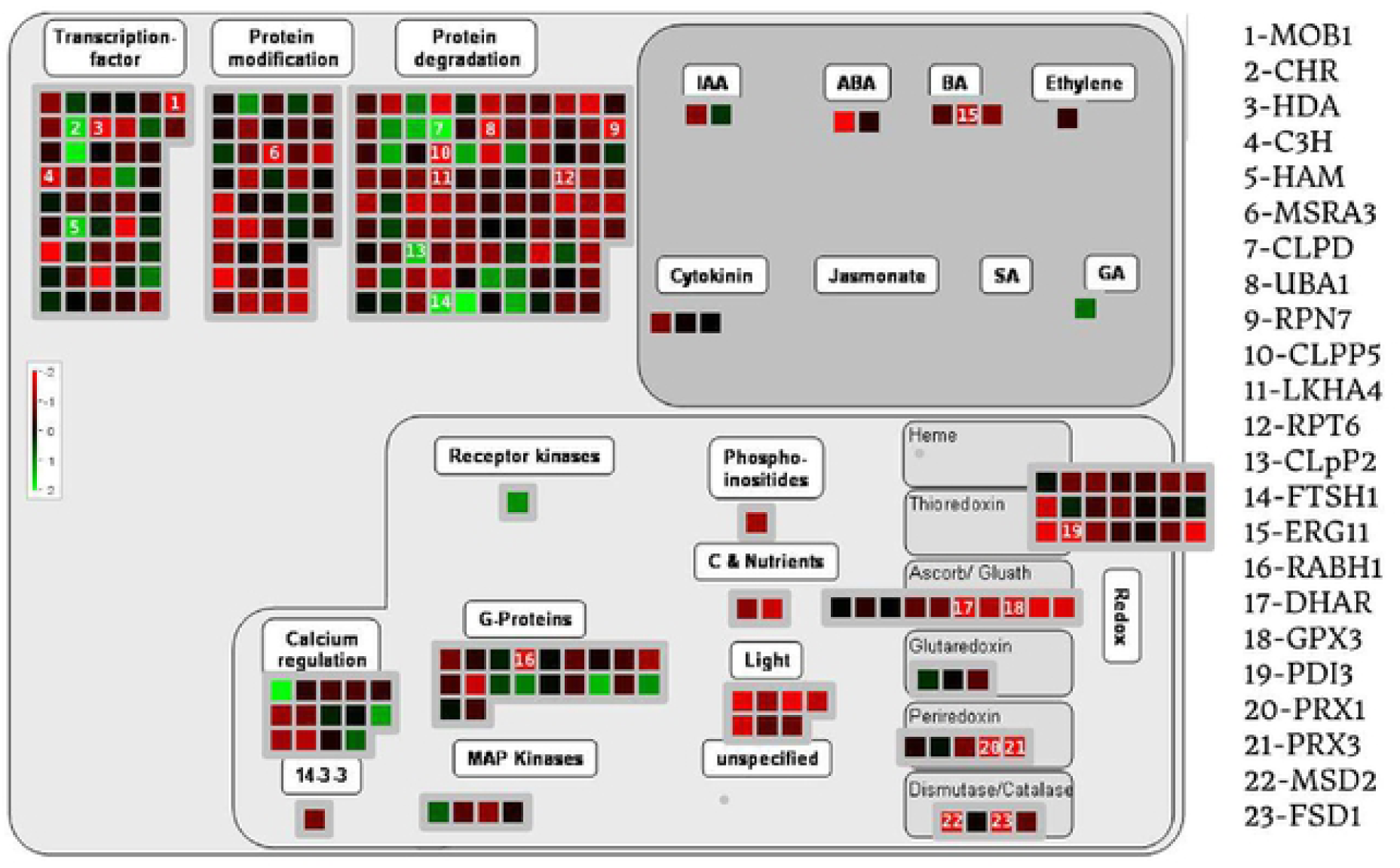
Regulation overview of diferentialy expressed genes under salt stress condition.

**Fig. 6:**
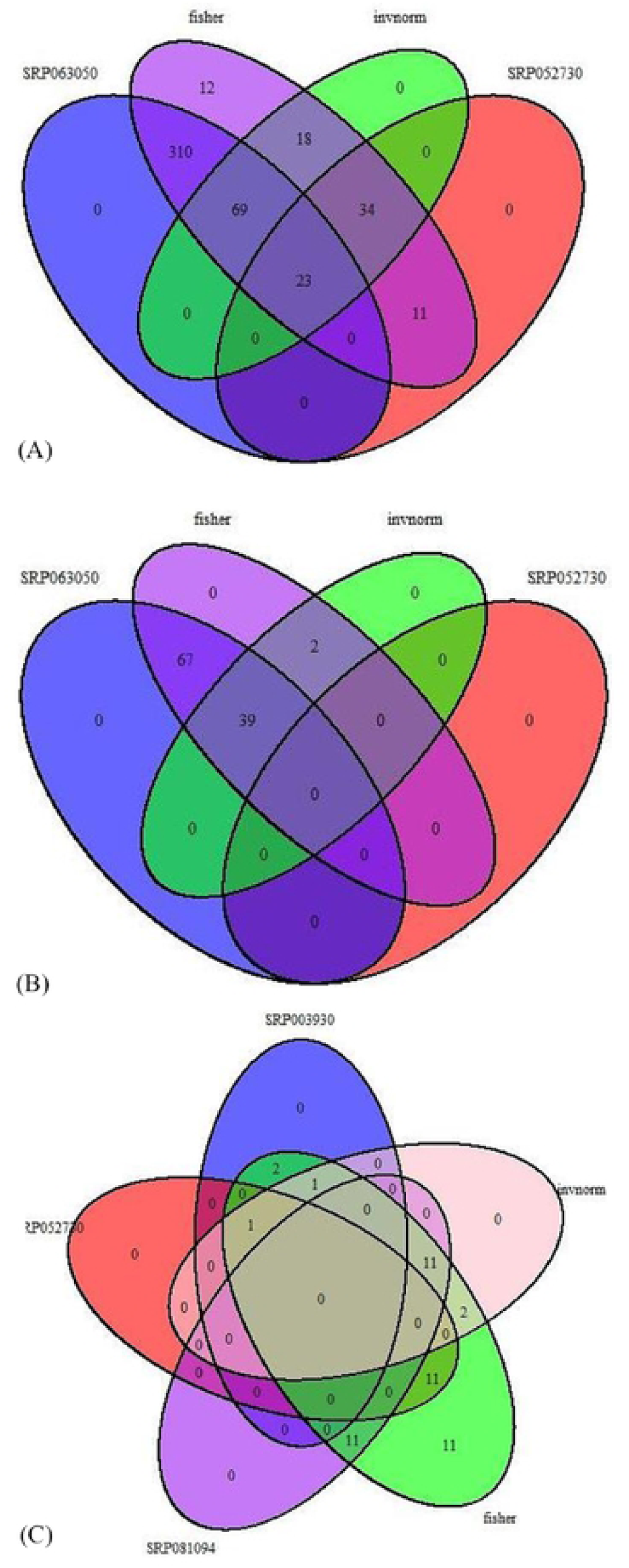
Venn diagram for DEGs of Salt (A), Light (B) and Nitrogen (C) stress (A) and meta results for each corresponding stress.

**Fig. 7:**
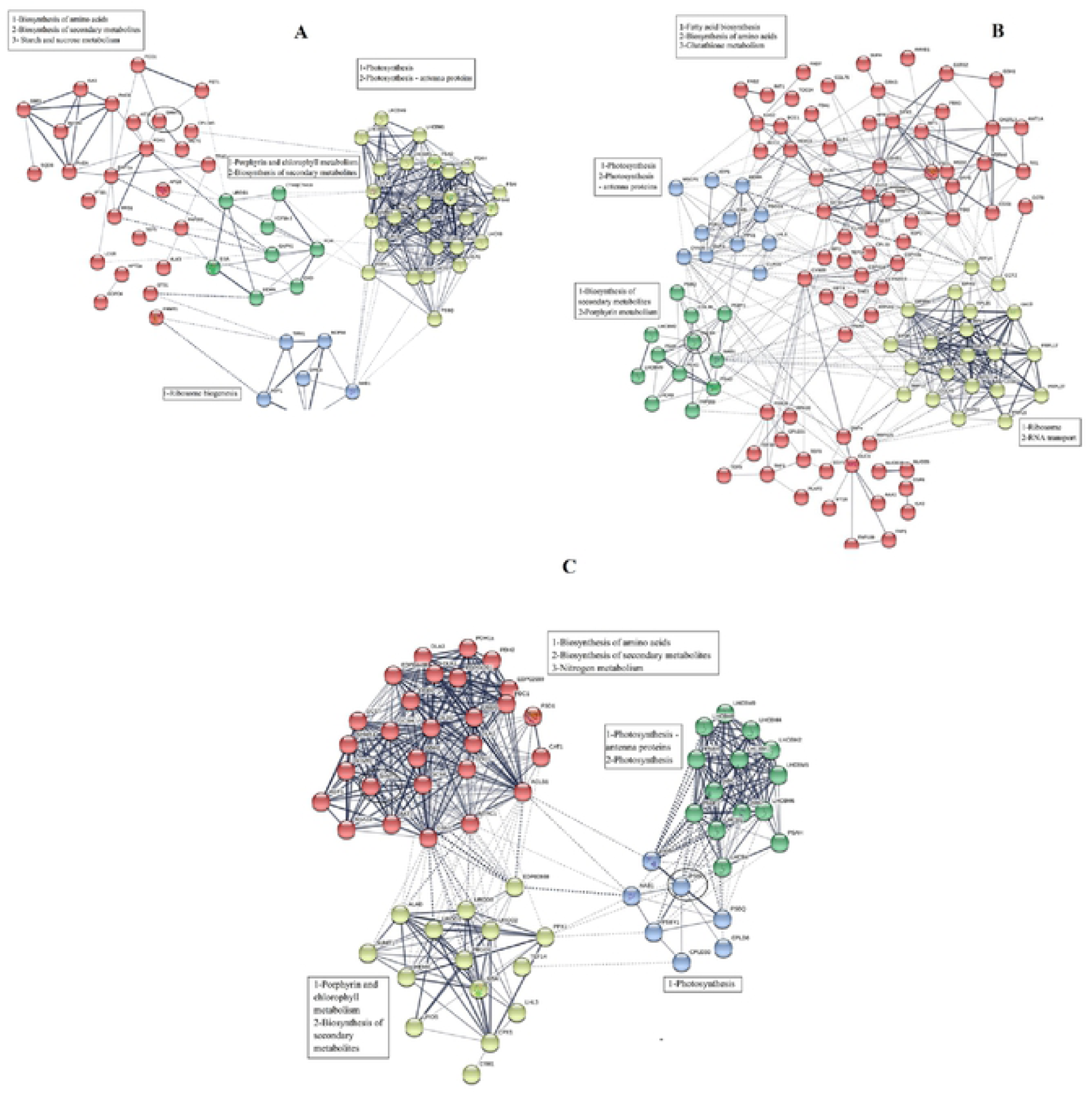
Protein-Protein interaction network of meta genes for light stress (A), Salt stress (B) and nitrogen deficiency (C) conditions in *D. tertio/ecta* based on the *chlamydomonas reinhardti* knowledge.

The module enrichment of the PPI networks also highlighted the cross talk between photosynthesis, fatty acids, starch, and sucrose metabolism under multiple stress conditions (Fig. 4). The KEGG enrichment of the identified meta-genes under the above-mentioned stress conditions suggested that tetrapyrrole biosynthesis was a key stress responsive pathway in *D. tertiolecta*. Among different genes involved in thetetrapyrrole biosynthetic pathways, glutamyl-tRNA reductase (*GluTR*), uroporphyrinogen decarboxylase (*UROD*), porphobilinogen synthase (*ALAD*), chlorophyllide a oxygenase (*CAO*), Mg^2+^-protoporphyrin IX monomethyl ester cyclase (*CHLE*), glutamyl-tRNA synthetase (*EARS*), and protochlorophyllide reductase (*POR*) were defined as meta-genes under high light condition (Supplementary Table S2); however, protoporphyrin/coproporphyrin ferrochelatase (hemH), geranylgeranyl diphosphate (chlP), hydroxymethylbilane synthase (*HMBS*), and GluTR were detected as meta-genes under high salt stress condition. Under nitrogen deficiency condition, coproporphyrinogen III oxidase (*CPOX*), geranylgeranyl diphosphate (chlP), and hydroxymethylbilane synthase (*HMBS*) were identified as meta-genes (Supplementary Table S2). Two main functions of the tetrapyrrole biosynthesis pathway are the chlorophyll (Chl) biosynthesis and plastid to nucleus retrograde signaling (Kobayashi and Masuda 2016). In contrast with previous findings in plants, our analysis indicated that a majority of meta-genes involved in the tetrapyrrole pathway genes are dominantly up-regulated by high light (Cihlář *et al*. 2016). In line with the results of our analysis, the up-regulation of tetrapyrrole biosynthesis genes under photo-oxidative stress condition has been documented, as reported in *Chlamydomonas reinhardtii* (Nogaj *et al*. 2005). It has been also reported that oxidative stress triggers chloroplast to nucleus communication (Wang *et al*. 2018). Similar expression patterns of the identified meta-genes were noticed under nitrogen deficiency condition. The coordinated expression of the tetrapyrrole intermediated with starch biosynthesis involved meta-genes was observed in this study. Moreover, the co-occurrence and physical interaction of the starch biosynthesis with tetrapyrrole biosynthesis involved meta-genes under high salt, high light, and nitrogen deficiency conditions confirm the hypothesis indicating that plastid to nucleus retrograde signals are essential for the expression of nuclear starch biosynthesis genes (Wang *et al*. 2018).

The meta-genes of the three abiotic stresses were also enriched in photosynthesis processes. The functional module enrichment of PPI highlighted this finding. Photosynthetic-reaction center complexes, light-harvesting complexes, and photosynthetic antennae were the most abundant functional groups of photosynthesis-involved meta-genes. Transcripts of light-harvesting chlorophyll (Chl) a/b-binding proteins B2 (Lhcb2) and *LHCB4* were constitutively decreased under the three abiotic stress conditions (Supplementary Table S2). *LHCs* allows the reaction centers of photosystem (PS) I and II to initiate charge separation and fuel the photosynthetic electron transport chain in the thylakoid membranes (Ishizaki et al., 2012; Perrine et al., 2012). It has been proposed that feedback regulation of Lhcb is essential to reduce the effects of reactive oxygen species on Chls and to accumulate metabolic intermediates under oxidative stress conditions (Wang *et al*. 2018). Hence, the reduced accumulation of the Mg-Proto, corresponded with decreased expression of *LHCB2* and *LHCB4*, proposed the regulatory effects of tetrapyrrole biosynthesis intermediates on Chl and carotenoid pigment biosynthesis under multiple stress conditions.

Furthermore, the KEGG enrichment analysis of the meta-genes and functional modules of PPI networks indicated that vitamin B6 metabolism, methane metabolism, ribosome biogenesis, folate biosynthesis, and tetracycline biosynthesis responded to the different stresses (Fig. 4 and Supplementary Table S2). Vitamin B6 metabolism and methane metabolism were specifically enriched in light stress meta-genes (Supplementary Table S2). Indeed, the accumulation of the vitamin B6 confers oxidative stress tolerance (Raschke *et al*. 2011). It has been also reported that the vitamin B6 alleviates oxidative damage of Chl and carotenoid pigments intermediates (Havaux *et al*. 2009).

The findings also indicated that ribosome biogenesis specifically enriched under salt stress (Fig. 4). Among protein synthesis apparatus involved genes in *D. tertiolecta*, nine large (*RPL15, RPL37, RPL27A, RPL8, RPL23A, MRPL9, MRPL17, MRPL27*, and *RPL7*) and 3 small (*RPSA, RPS19, RPS27*) subunits of ribosomal proteins were defined as core responsive-genes under salt stress condition. Most interestingly the cytosolic ribosomal proteins, including the *RPL15, RPL23A, RPL27A, RPL37, RPL7, RPL8, RPS19*, and RPS27 were found to be up-regulated under salt stress condition, while mitochondrial ribosomal proteins such as *MRPL9, MRPL17, MRPL27*, and *RPSA* were down-regulated under salt stress (Supplementary Table S2). The salt stress-dependent down-regulation of *MRPL9, MRPL17*, and *MRPL27* may contribute to mitochondrial functionality through maintaining redox status. It was proposed that the transcriptional control of the ribosome synthesis is an important regulatory point of signal and regulon generation under salt stress condition (Omidbakhshfard *et al*. 2012). Although, microalgae ribosomal proteins have not yet been studied in detail, preliminary studies showed that a mutation in the RP genes (*RPS18A, RPL24B, RPS5B, RPS13B*, and *RPL27A*) leads to decreased cell division and growth rate (Liang *et al*. 2017).

## Conclusion

The present study employed a meta-analysis of RNAseq studies under the abiotic stress conditions, including high light, high salt, and nitrogen deficiency on *D. tertiolecta*. This approach magnifies the efficiencies in the identification of key and core stress response genes/pathways, which might be overlooked by previous studies. The comparative analysis of differential expression analysis and KEGG pathway enrichment of the three stress conditions indicated the existence of shared and unique pathways under high light, high salt, and nitrogen deficient conditions. The shared stress-responsive genes were found to be involved in the tetrapyrrole biosynthesis pathway, starch, and sucrose metabolism. These changes in metabolism allow *D. tertiolecta* to maintain its homeostatic state and redirect the carbon flux to over-accumulate multiple secondary metabolites.

## Author contributions

Conceived and designed the experiments: BP. Analyse the data: BP, MF, writing the paper: BP, SAM, and MAH.

## Acknowledgements

The authors are grateful to Dr Anusuya Willis (CSIRO, Australia) for critical reviewing of manuscript. The financial support of the Agricultural Biotechnology Research Institute of Iran (ABRII) is gratefully acknowledged.

---

**Table 1**. **Raw data set ID, run accession and read count (before and after trimming)**.

**Fig. 1: Metabolic overview of differentially expressed genes in Light stress conditions in *D. tertiolecta*. Green and red box represented the down and up-regulation in corresponding stress, respectively. Up-regulation of myo-inosytol-phosohate synthase, phospholipid synthase PGP1b and fatty acid desaturase are indicated. Most of the genes involved in photorespiration were down-regulated in response to light stress**.

**Fig. 2: Regulation overview of differentially expressed genes under salt stress condition. Most of the genes involved in protein modification and degradation were down regulated in response to salt stress**.

**Fig. 3: Venn diagram for DEGs of Salt (A), Light (B) and Nitrogen (C) stress (A) and meta-analysis results for each corresponding stress. In total 621, 65 and 149 meta-genes were defined for salt, nitrogen deficiency and high light stress conditions, respectively. Most of the identified meta-genes were down regulated at all of the mentioned stresses**.

**Fig. 4: Protein-Protein interaction network of meta genes for light stress (A), Salt stress (b) and nitrogen deficiency (C) conditions in *D. tertiolecta* based on the *chlamydomonas reinhardti* knowledge. The results indicate that the key meta-analysis genes (signed with red circle) tend to be located in a hub situation of the stress-specific PPI networks**

